# Brassinosteroids and Flavonols Confer Temperature Stress Tolerance to Pollen Tube Germination and Growth

**DOI:** 10.1101/2024.10.27.620467

**Authors:** Kumi Matsuura-Tokita, Ayaka Sakai, Takamasa Suzuki, Akihiko Nakano, Tetsuya Higashiyama

## Abstract

Successful reproduction is a prerequisite for the maintenance of species in the era of climate change. In agriculture, temperature has a significant impact on pollen tube elongation and attraction, and consequently, on the efficiency of fruiting and crop yields. Brassinosteroids (BRs) are plant steroid hormones involved in pollen tube germination, elongation, and capacitation. To elucidate the effect of BR signaling in pistils during the reproductive process, transcriptome analysis of BR receptor (BRI1) mutant pistils was performed, which indicated that genes involved in flavonol biosynthesis were downregulated in *bri1* pistils. Flavonols are essential for pollen tube growth in some plants and facilitate pollen tube elongation under high-temperature conditions. To investigate the effects of BR and flavonols on pollen tubes under temperature stress, *in vitro* assays were performed using both the compounds. Brassinolide (BL), which is the most active BR, promoted pollen tube germination at high, low, and optimal temperatures. Germination was further enhanced by the flavonol, quercetin, in the presence of BL especially at low temperatures. *in vivo* analysis supported the observation that when BL and quercetin were supplied concurrently, the pollen tube guidance rate was rescued at low temperatures. Quercetin enhanced pollen tube elongation under all the conditions tested at an optimal concentration of 15 μM. BL elevated reactive oxygen species (ROS) under all temperature conditions, whereas quercetin regulated ROS synergistically with BR. Thus, BRs and flavonols supplied by pistils are suggested to function in pollen tube germination and elongation through the maintenance of ROS homeostasis.

## Introduction

Plants produce a wide variety of secondary metabolites and adapt to their surrounding environments. They serve as defense mechanisms against pathogens, protect against ultraviolet radiation, and provide relief from stressors such as drought, salt stress, and temperature shifts (Bowne et al., 2012; Ithal and Reddy, 2004; Ma et al., 2014; Muhlemann et al., 2018). Flavonols are secondary metabolites that protect plants from ROS generated by ultraviolet radiation. Additionally, they facilitate pollen tube germination and elongation (Mo et al., 1992; Ylstra et al., 1994, 1992). Flavonols are present in both male and female reproductive organs of *Petunia* and are essential for pollen tube germination (Ylstra et al., 1994) However, as the flavonol-synthesizing mutant (*tt4-1*) of *Arabidopsis thaliana* was fertile, it was postulated that flavonols were not essential for reproduction in *A. thaliana*.

We have previously shown that the plant steroid hormone brassinosteroid (BR) promotes pollen tube capacitation and the expression of genes involved in pollen tube attraction in the ovules (Matsuura-Tokita et al., 2024). BRs are important in reproductive development in plants (Lima and Figueiredo, 2024). To further investigate the signaling mechanisms of BR within the pistil, we performed a transcriptome analysis of the *bri1* mutant pistil. The expression of transcription factors and enzymes involved in the biosynthesis of the secondary metabolite, flavonol, was downregulated in the *bri1* mutant pistil compared to that of the wild type. Flavonols act as ROS scavengers to reduce ROS under high temperatures in the tomato pollen tubes (Muhlemann et al., 2018). In this study, we examined the role of brassinolide (BL) and a flavonol, quercetin, in pollen tube germination and elongation *in vitro* under temperature stress. BL promoted pollen tube germination, presumably through the production of ROS. Quercetin demonstrated a reduction in ROS at both elevated and optimal temperatures, although quercetin in combination with BL did not show a reduction in ROS at low temperatures, leading to ROS levels comparable to the optimal temperature. By the application of BL and quercetin, *in vivo* analysis indicated the rescue of pollen tube guidance ability at low temperatures. These findings indicate that BL and quercetin function synergistically to maintain ROS homeostasis in pollen tubes, thereby conferring temperature-stress tolerance.

## Results

### Gene expression involved in flavonol synthesis was decreased in *bri1* pistils

In a previous study, pollen tubes elongated through *bri1* pistils showed decreased guidance ability in semi-*in vivo* assay (Matsuura-Tokita et al. 2024). Transcriptome analysis of wild type and *bri1* pistils (Supplementary Data Set 1) was performed to elucidate the underlying mechanisms responsible for the observed differences in pollen tube activity. KEGG pathway analysis revealed that genes involved in the “flavonoid biosynthesis” pathway showed decreased expression in *bri1* pistils (logFC < -1, FDR < 0.05, >1 rpm) (Fig. 1A) (Supplementary Data Set 2). We subsequently investigated alterations in the expression of three transcription factors involved in flavonol synthesis: *MYB11, 12*, and *111* (Fig. 1B–D) and three biosynthetic enzymes, *CHS, F3H*, and *FLS* (Fig. 1E–G). Consequently, the expression of all factors was reduced in *bri1* pistils compared to wild-type plants. These genes, except *MYB12*, were highly expressed in the pistils (*Arabidopsis* eFP Browser, http://bar.utoronto.ca/efp/cgi-bin/efpWeb.cgi) (Supplementary Fig. S1). In this study, preliminary experiments were conducted with three flavonols that promote tobacco pollen tube germination (Ylstra et al., 1992), namely, quercetin, myricetin and kaempferol. We focused on quercetin, which was the most effective (data not shown). Quercetin is more active than myricetin and kaempferol in immature pollen experiments in tobacco (Ylstra et al., 1992).

**Figure 1.**
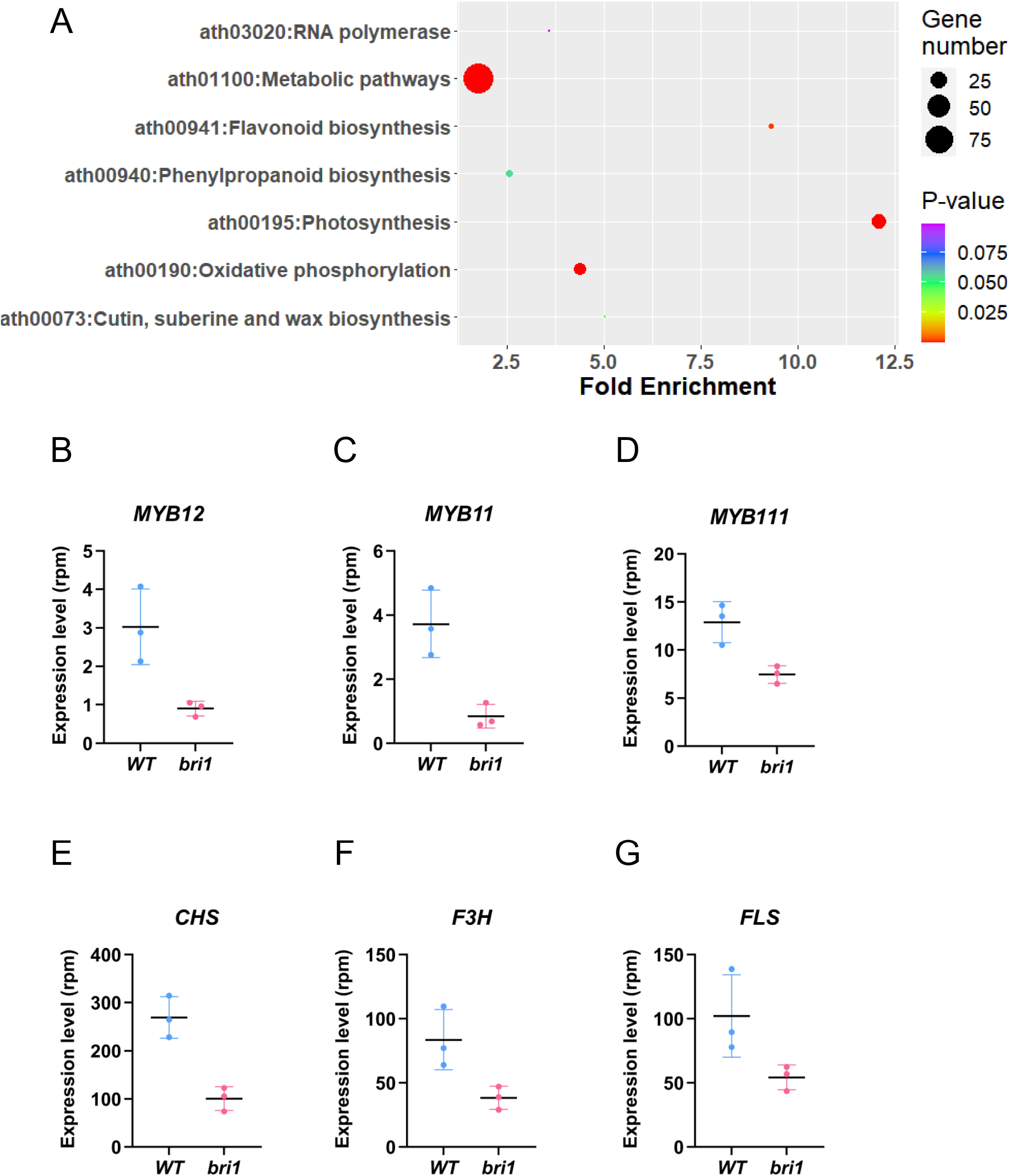
Expression of flavonol biosynthesis genes was downregulated in *bri1* pistils. A. KEGG pathway analysis of genes downregulated in *bri1* pistils. Genes involved in “Flavonoid biosynthesis” showed a high Fold Enrichment. B-G. Comparison of gene expression related to flavonol biosynthesis between wild type and *bri1* pistils. B-D. Transcription factors E–G. Enzymes involved in flavonol synthesis. B. MYB11, C. MYB12, D. MYB111, E. CHS, F. F3H, and G FLS.

In previous studies, a mutant (*tt4-1*) defective in chalcone synthase (*CHS*), a key enzyme for flavonol biosynthesis in *A. thaliana* was fertile (Burbulis et al., 1996; Ylstra et al., 1996), leading to the recognition that flavonols are not essential for reproduction in *Arabidopsis*. We quantified the seed number of the self-pollinated *myb11/12/111* triple mutant, which is also deficient in flavonol synthesis (Stracke et al., 2007), and found a significant reduction in seed number per silique (Supplementary Fig. S2). The data suggests that *myb11/12/111* triple mutant deficient in flavonol synthesis is not infertile but defective in reproduction.

### BL promoted pollen tube germination under all the temperature conditions tested

The germination rate and length of pollen tubes were subsequently quantified under different temperature conditions of 15°C (low), 23°C (optimal), and 30°C (high). Pollen tube elongation medium was prepared with BL concentration of 10 μM, according to a previous report (Vogler et al., 2014). Quercetin was applied at final concentrations of 0, 5, 10, 15, and 20 μM respectively. BL promoted the germination of pollen tubes under all the temperature conditions tested (Fig. 2 A-C). Quercetin alone also promoted the germination rate in a dose-dependent manner at 23 °C. When applied with BL, the germination rate increased. Under high-temperature conditions (30°C), the average germination rate was approximately 70% in the absence of any compounds (Fig. 2 C). The germination rate increased to approximately 90% when BL and quercetin were simultaneously applied. No dose-dependent effects of quercetin were observed at 30°C.

**Figure 2.**
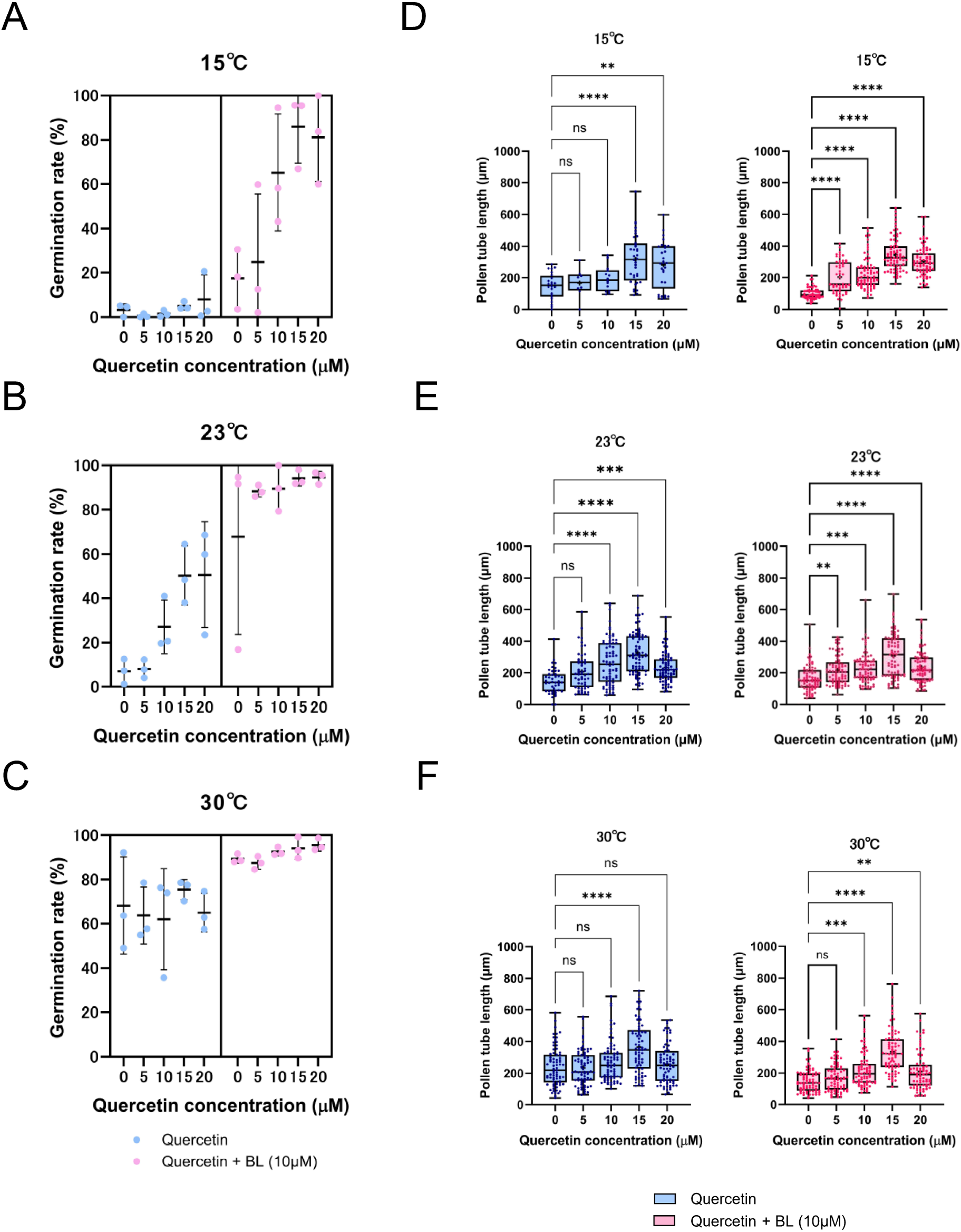
Pollen tube germination rate and length under different temperature conditions with BL and quercetin. The BL concentration was maintained at 0 or10 μM, while quercetin concentrations varied from 0 to 20 μM, with increments of 5 μM. A-C. Pollen tube germination rates at 15°C (A), 23°C (B), and 30°C (C). Mean (SD). D-F. Pollen tube length at 15°C (D), 23°C (E), and 30°C (F). Kruskal Wallis test was performed to determine significant differences among groups. Dunn’s test was performed to determine pairwise comparisons among all groups.

Under low temperature conditions (15°C), quercetin alone did little to promote germination at any concentration. However, when quercetin and BL were added concomitantly, germination was promoted in a dose-dependent manner by quercetin, and the maximum germination rate was observed when 15 μM quercetin was added. Under very low-temperature conditions (4°C), germination occurred only when BL was added (Supplementary Fig. S3). No dose-dependent effects were observed for quercetin. Therefore, the contribution of BL to pollen tube germination under low-temperature conditions is prominent. The dose-dependent effect of quercetin on the germination rate in the presence of BL at 15°C suggests the synergistic effects of both BL and quercetin on pollen tube germination.

### Quercetin promoted pollen tube elongation under high and low temperature conditions

At 15, 23, and 30°C, pollen tube elongation was enhanced by quercetin in a dose-dependent manner, both in the presence and absence of BL, and the optimal concentration of quercetin was 15 μM (Fig. 2D-F). At 4°C, when BL and quercetin were applied concurrently, pollen tube length exhibited a significant increase in the presence of quercetin at the optimal concentration of 15 μM (Supplementary Fig. S3B). Although quercetin alone showed no positive effect on pollen tube length, synergistic effects were prominent under low-temperature conditions.

### ROS concentration in pollen tubes was elevated by the application of BL

Previous studies have reported that in tomatoes, pollen tube germination and elongation are impaired in mutants exhibiting reduced flavonol content in the pollen (Muhlemann et al., 2018). In this mutant, reactive oxygen species (ROS) accumulate to a greater extent than in the wild type under normal temperature conditions, suggesting that flavonol functions as a scavenger of ROS. Consequently, we aimed to compare ROS signals in pollen tubes in the presence of exogenous 10 μM BL and/or 15 μM quercetin at low (15°C), optimal (23°C), and high temperatures (30°C). The average fluorescence intensity in the apical region of pollen tubes (100 μm from the tip), was quantified. ROS staining by CM-H_2_DCFDA (Thermo Fischer) at 23°C showed that pollen tube ROS levels were higher when BL was applied. Quercetin-only treatment resulted in lower ROS levels than the negative control. When BL and quercetin were applied concurrently, ROS levels were comparable to those in the control.

At 30°C, the ROS signal in pollen tubes was high compared to that at other temperatures. The exposure time of the imaging conditions was reduced to one-fifth of that at 23°C and 15°C to prevent saturation of fluorescence. At 30°C and 23°C, the pollen tube ROS signal was high when BL was applied. Quercetin lowered the ROS levels in the presence and absence of BL (Figure 3). Conversely, at 15°C, the pollen tube ROS level was elevated when BL and quercetin were applied concurrently compared to the application of BL alone.

**Figure 3.**
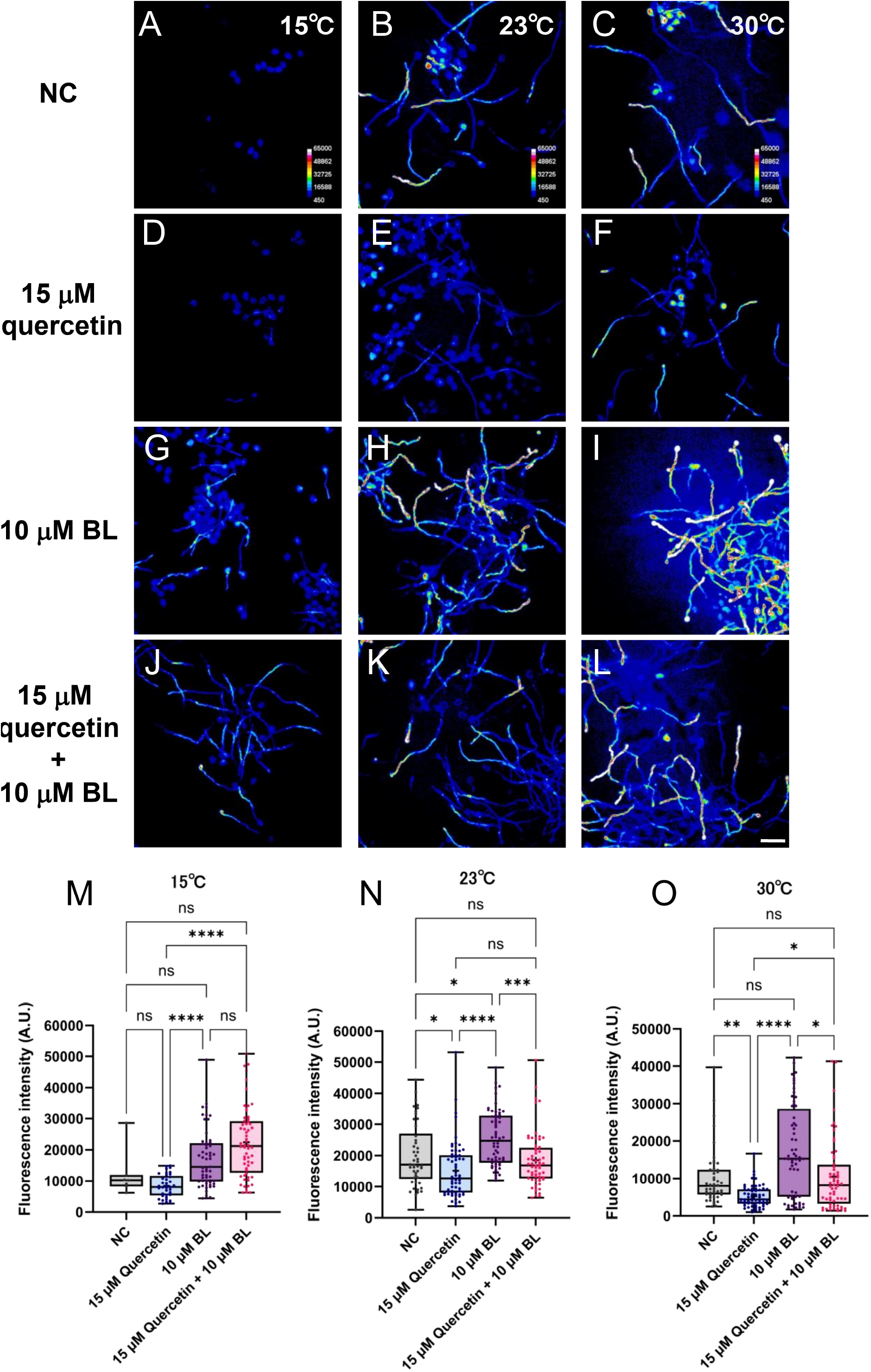
ROS staining in pollen tubes under different temperature conditions with BL and quercetin. A–C. *in vitro* cultured pollen tubes without any compounds (NC) at 15°C, 23°C, and 30°C, respectively. D–F. *in vitro* cultured pollen tubes incubated with 15 μM quercetin at 15°C, 23°C, and 30°C, respectively. G–I. *in vitro* cultured pollen tubes incubated with 10 μM BL at 15°C, 23°C, and 30°C, respectively. J–L. *in vitro* cultured pollen tubes incubated with 15 μM quercetin and 10 μM BL at 15, 23, and 30°C, respectively. Pollen tubes were stained with CM-H_2_DCFDA for 20 min. Scale bar, 50 μm. M–O. Quantification of ROS fluorescence intensity of pollen tube tip region (100 μm from the tip) at 15°C, 23°C, and 30°C, respectively.

### *in vivo* effects of BL and quercetin at low temperatures

To examine *in vivo* effects of BL and quercetin under low-temperature conditions, wild type plants were subjected to pollination at 15 °C. Emasculated pistils were supplemented with a solution containing 10 μM BL and 15 μM quercetin followed by manual pollination. Following overnight incubation of the plants at 15°C, pistils were harvested, and aniline blue staining was performed to visualize pollen tube growth within the pistils. Pistils treated with BL and quercetin exhibited a higher rate of pollen tube attraction at 15°C than the negative control (Figure 4A).

**Figure 4.**
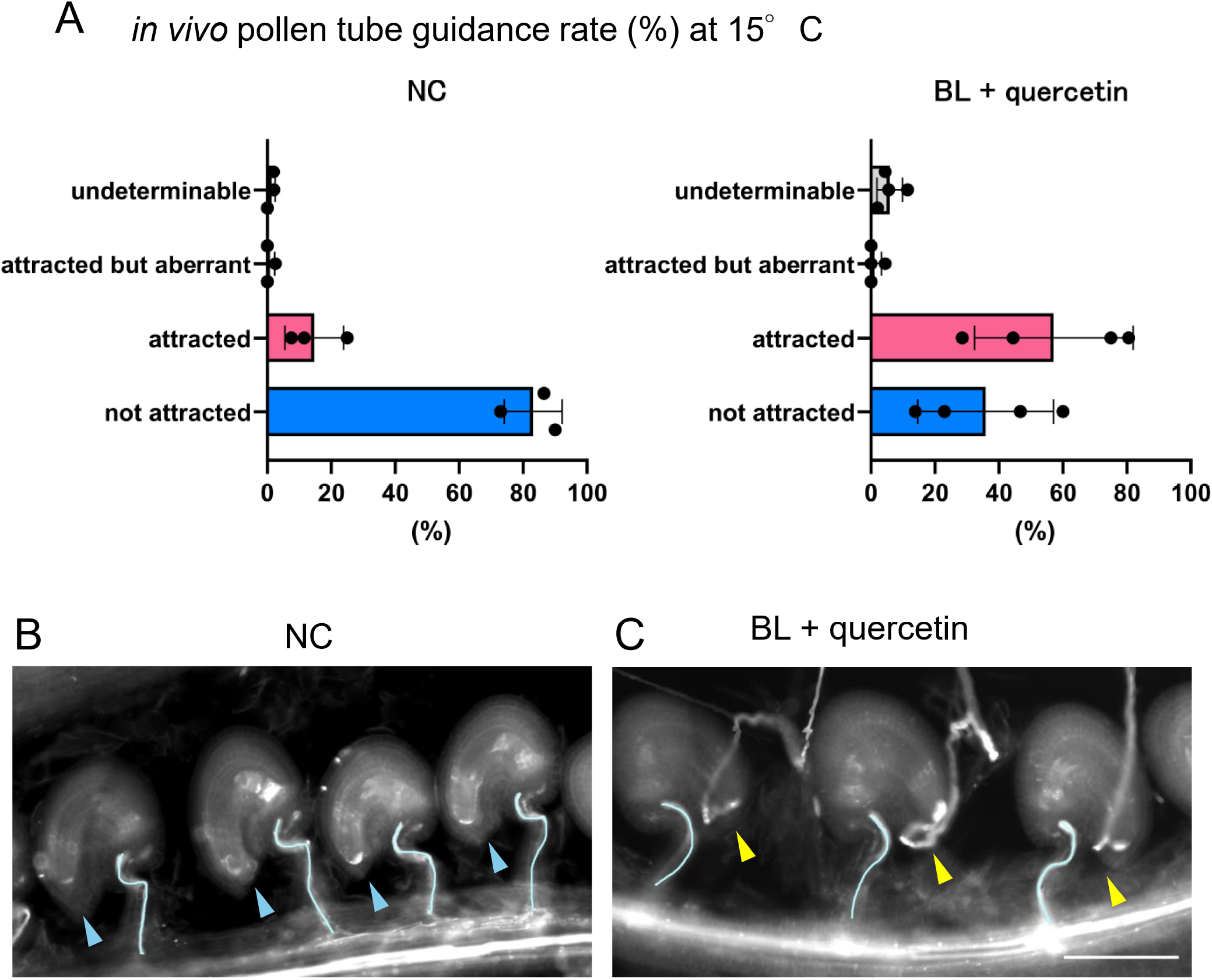
*in vivo* pollen tube guidance at 15°C. Pistils treated with solvents (NC) or with BL and quercetin (BL + quercetin) were hand pollinated and incubated for 24h at 15°C. A. Percentage of attracted and unattracted ovules under each condition. B. Aniline blue staining of pistils treated with solvents (NC). Blue arrowheads indicate the micropylar regions of non-attracted ovules. C. Aniline blue staining of pistils treated with BL and quercetin. The yellow arrowheads indicate the micropylar regions of the attracted ovules. Scale bar, 100 μm. Blue lines in B and C indicate vascular bundles.

## Discussion

In the present study, we showed flavonol synthesis was regulated downstream of BRI1 signaling pathway in pistils. Application of BL and a flavonol, quercetin, significantly enhanced *Arabidopsis* pollen tube germination and elongation under both high- and low-temperature stress conditions. Our study indicated that BRs and flavonols supplied by pistils confer temperature stress tolerance by maintaining ROS homeostasis in pollen tubes.

### Pollen tube germination and BL

Although quercetin had a modest influence on pollen tube germination, the effect of BL was significant. The addition of BL under all temperature conditions increased ROS signals at the pollen tube tip, suggesting that BL promotes germination by stimulating ROS synthesis. ROS are important for pollen tube germination in plants (Bowne et al., 2012; Speranza et al., 2012). Nevertheless, transcriptome analysis of pollen tubes treated with BL (Matsuura-Tokita et al., 2024) revealed that the expression of RBOHH and RBOHJ, which synthesize ROS in pollen tubes, remained unaffected. Similarly, the expression of receptor complexes LLG2/3, BUPS1/2, and ANX1/2, which function upstream of RBOHH/RBOHJ (Feng et al., 2019; Zhang et al., 2020), was not upregulated. On the other hand, a series of factors that regulate intracellular ion gradients, including the cation/H+ exchanger (CHX) family, AHA4, NHX4, and AHA6, are upregulated, which may induce ROS production (Esparza-Reynoso et al., 2023). The expression of calcium-related genes such as CNGC7, ROP4/8, and ROPGEF11/13 was also increased. As the Ca^2+^-binding EF-hand motifs of RBOH/RBOHJ(Kaya et al., 2014) are essential for ROS production, it is possible that BL indirectly increases ROS through these factors.

### Function of flavonols at low temperatures

At high temperatures, ROS signals were reduced by quercetin. At low temperatures, ROS signals was not reduced by quercetin in the presence of BL. Quercetin has been reported to be a ROS scavenger, however, in this study we hypothesized that there must be other functions under low temperature conditions. ROS signal intensity in the presence of both BL and quercetin was comparable at 15°C and 23°C, suggesting that these compounds work synergistically to maintain ROS homeostasis in pollen tubes. Under all experimental conditions examined, pollen tube lengths varied in a quercetin concentration-dependent manner, both in the presence and absence of BL, with the maximum length at a quercetin concentration of 15 μM. This is the first report indicating the promotive effects of exogenously applicated quercetin on pollen tube elongation and germination under temperature stress. However, the reception and signaling mechanisms of flavonols are largely unknown. Further investigation is necessary to elucidate the underlying molecular mechanisms.

### *in vivo* effects of BL and quercetin

The pollen tube guidance rate was increased by BL and quercetin under low-temperature conditions *in vivo*. The data in Figure 4 clearly indicate that exogenous application of these compounds was useful for increasing fertilization under temperature stress conditions *in vivo*. In future studies, it is expected that these effects will be verified in plants other than *Arabidopsis*, especially in crops and fruits. Utilizing biostimulants, such as BRs and flavonols, for improved plant reproduction in severe environments will open up new ways toward sustainable agriculture.

### Materials and methods Plants and culture conditions

*Arabidopsis thaliana* was used in this study. Ecotype Columbia (*Col-0*) was used as the wild type. CS860325 (SALK_ 041648) was purchased from ABRC and was used as *bri1-10* (Vogler et al., 2014). *myb11/12/111* triple mutant (N9815) was purchased from ABRC.

The plants were grown under conditions of 16 h light–8 h dark, 22°C, and approximately 30 % humidity. A solution of Hyponex 6-10-5 (Hyponex Japan) diluted 1 per 5000 was added to water as needed.

### RNAseq analysis of pistils

Mature pistils were collected 24h after emasculation (four pistils for wild-type, six pistils for *bri*1*-10*, three replicates each). mRNA was extracted and sequenced as previously described (Matsuura-Tokita et al. 2024). Briefly, mRNA was extracted using the Dynabeads mRNA DIRECT Micro Kit (Thermo Fischer). A library was constructed using the Illumina TruSeq RNA Sample Preparation Kit ver. 2. Sequencing was performed using NextSeq500 (Illumina) and NextSeq500 High Output Kit v2.5 (Illumina). Raw reads with adapter sequences were trimmed using bcl2fastq (Illumina). Low-quality nucleotides (QV < 25) were masked by N and short reads (<50bp) were discarded using the original script.

Using Bowtie, the remaining reads were mapped to the cDNA reference sequence, with the following parameters: ‘--all –best --strata’ (Langmead et al., 2009). Subsequently, reads were counted by transcript models. The data were analyzed using R 4.0.2 statistical software. Differentially expressed genes were identified by the edgeR software (Robinson et al., 2010). KEGG pathway analysis was conducted by The Database for Annotation, Visualization and Integrated Discovery (DAVID) 2021 (Huang et al., 2009; Sherman et al., 2022)

### *In vitro* observation of pollen tubes

The medium for pollen tube germination and elongation was prepared as described (Boavida and McCormick, 2007; Muro et al., 2018). Epibrassinolide (SIGMA-Aldrich) was dissolved in 90 % ethanol at the concentration of 10 mM and quercetin (SIGMA-Aldrich) was dissolved in DMSO at the concentration of 10 mM.

The flowers were collected, and the stamens were cut out using tweezers under a stereomicroscope. The anthers were then placed on the medium to distribute pollen grains. Pollen grains from three flowers were used per sample. Pollen tubes were incubated overnight in the dark at 23°C. 20 μl drops of 0.1% (w/v) aniline blue in a 2% K_3_PO_4_ solution were used for pollen tube observation. An upright microscope (ZEISS Axio Imager2) was used for observation. Quantitative experiments were performed three times. Based on these images, the pollen tube length and pollen germination rate were calculated.

### Image analysis

Images of pollen tubes grown *in vitro* stained with aniline blue and DIC images were captured using an upright microscope (ZEISS Axio Imager2). Based on these images, the pollen tube length and pollen germination rate were quantified. For germination rate, depending on previous report examining the effect of flavonol on Tobacco (Ylstra et al., 1992), germination was defined as pollen tube length longer than half the size of the pollen grain. Clusters with more than 20 pollen grains in contact with each other were not included in the measurement because it was difficult to determine whether they had germinated. Pollen tube lengths were measured for 25 pollen tubes per sample.

### ROS staining of pollen tubes

Pollen grains from three flowers were distributed onto pollen tube germination medium and incubated for 4 hours under different temperature conditions. 10 μl of 5 μM CM-H_2_DCFDA (Thermo Fischer), a fluorescent reagent for detecting ROS, was dropped onto the medium and allowed to stand for 20 minutes each for observation. For the quantification of ROS, the average fluorescence intensity of the tip region (100 μm from the tip) was measured. Sixty pollen tubes were collected for each condition. Confocal microscopy was performed using a spinning-disk confocal scanner unit (CSU-X1; Yokogawa Electric).

### *In vivo* observation of pollen tubes in pistils

Emasculation was performed 24h before pollination. Wild type plants were moved to 15°C from 23°C, 2h prior to pollination. Emasculated pistils were dipped in 5% ethanol with 10μM BL and 15 μM quercetin, or 5% ethanol with solvent as the negative control. Subsequently, hand pollination was performed and the plants were incubated for 24h at 15°C. Pistils were collected, fixed, and stained with aniline blue as described previously (Ishiguro et al., 2001).

Pistils were fixed overnight in a solution of ethanol/acetic acid (3:1) at room temperature. They were softened by 4M NaOH overnight, washed three times with distilled water, and stained with 0.1% (w/v) aniline blue in 2% K_3_PO_4_ solution for 2h. Images were taken by inverted microscope (Nikon ECLIPSE T*i*) equipped with ORCA-Fusion BT Digital CMOS camera (Hamamatsu Photonics). Attracted/non-attracted ovules were counted using the Fiji software (ImageJ 1.52p, Java1.8.0_172 (64 bit)) (Schindelin et al., 2012). Three pistils were subjected for analysis as negative control, four pistils were used as BL+Q.

## Funding

This work was supported by the Japan Society for the Promotion of Science (JP15J40125 to K.M., JP18K19333 to T.H., JP21K18235 to T.H. and K.M., 22H04980 to T.H., and 22K21352 to T.H.). and CREST, Japan Science and Technology Agency (JPMJCR20E5 to T.H.).

## Acknowledgments

We would like to thank Yoshikatsu Sato, Masahiro Kanaoka, Narie Sasaki, Yusuke Kimata, Minako Ueda for technical support; Ayami Furuta for technical assistance.

## Author Contributions

**K.M. and T.H. designed research; K.M**., **A.S**., **and T.S. performed research; K.M**., **A.S**., **T.S. analyzed data; and K.M**., **A.S**., **T.S**., **A.N**., **and T.H. wrote the paper**.

## Disclosures

### Competing Interest Statement

The authors declare no conflict of interest.

## Notes

### Competing Interest Statement

The authors have declared no competing interest.

