## Supplementary information for "Brassinosteroids and Flavonols Confer Temperature Stress Tolerance to Pollen Tube Germination and Growth"

**Arabidopsis eFP Browser**

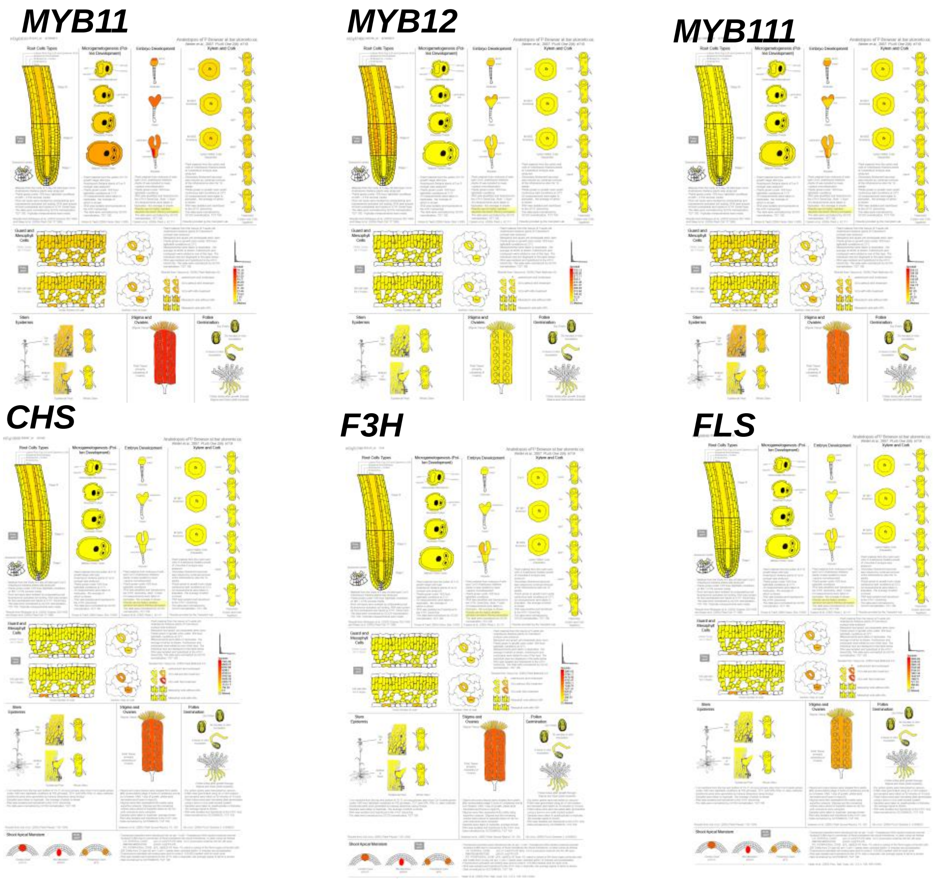


**Supplementary Fig. S1. Expression pattern of genes related to flavonol synthesis.** Tissue-specific expression of MYB11, MYB12, MYB111, CHS, F3H, and FLS in Arabidopsis eFP browser (Winter et al., 2007)**.**

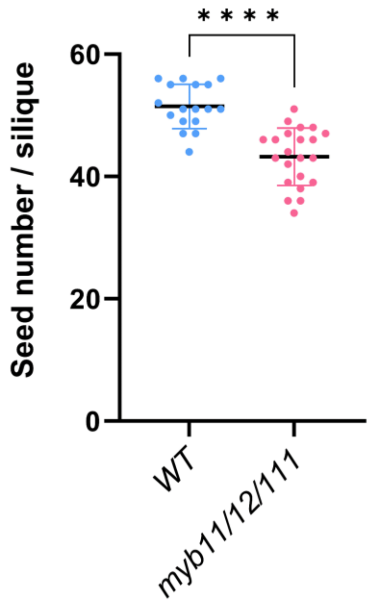


**Supplementary Fig. S2. Comparison of seed number per silique of wild type and *myb11/12/111* triple mutants.** The Mann-Whitney U test ($p<0.05$) was used for statistical analysis. Seventeen siliques of wild-type and 22 siliques of myb11/12/111 were quantified.

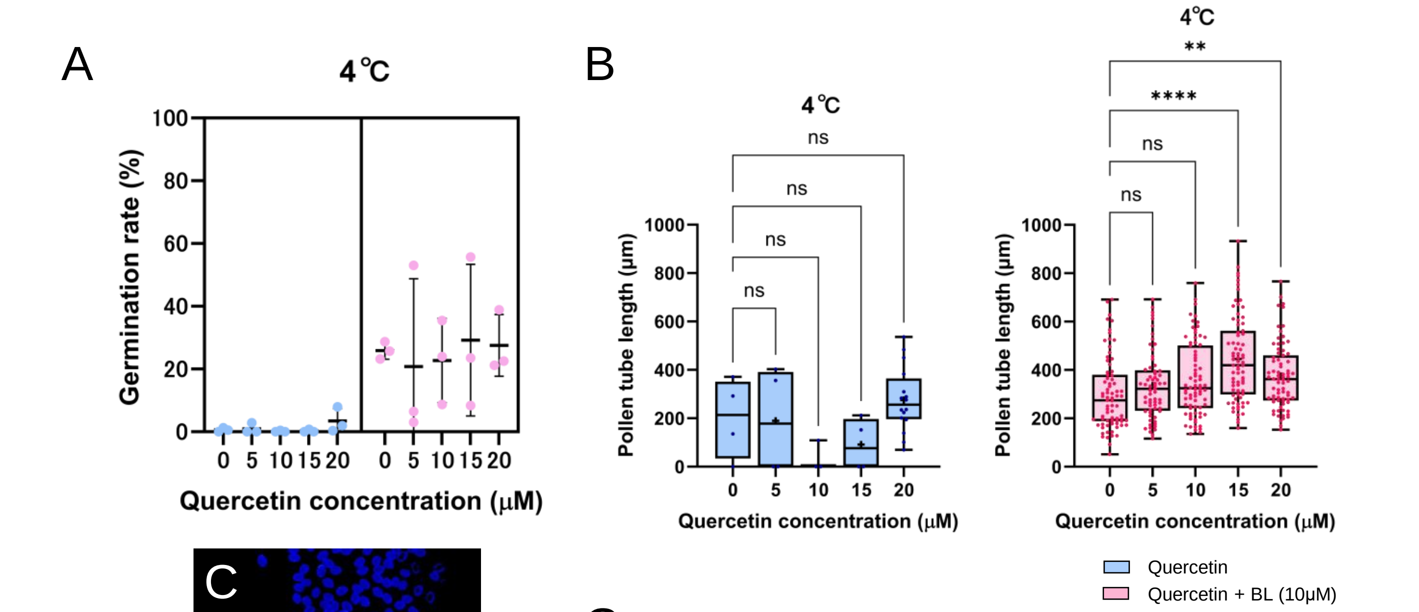


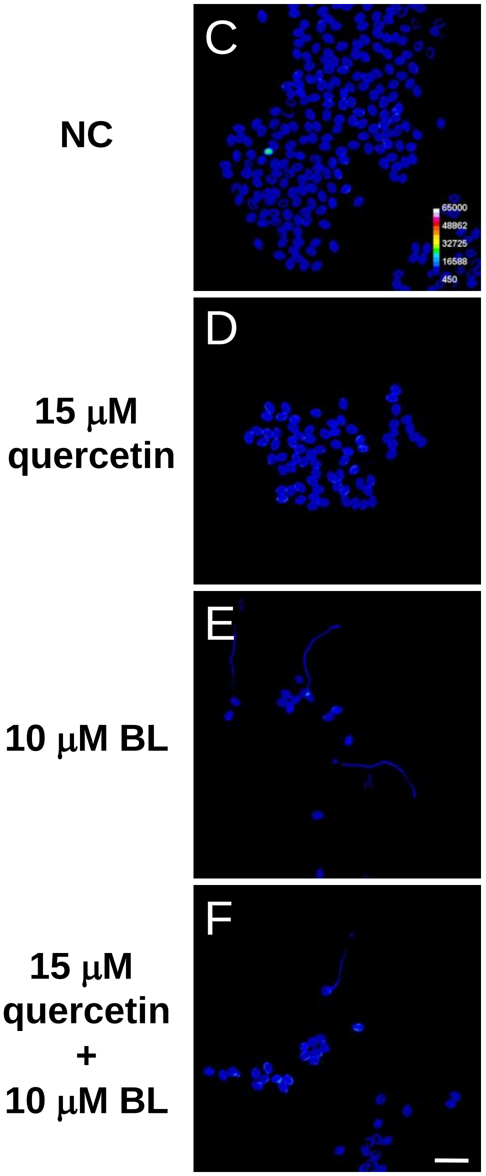


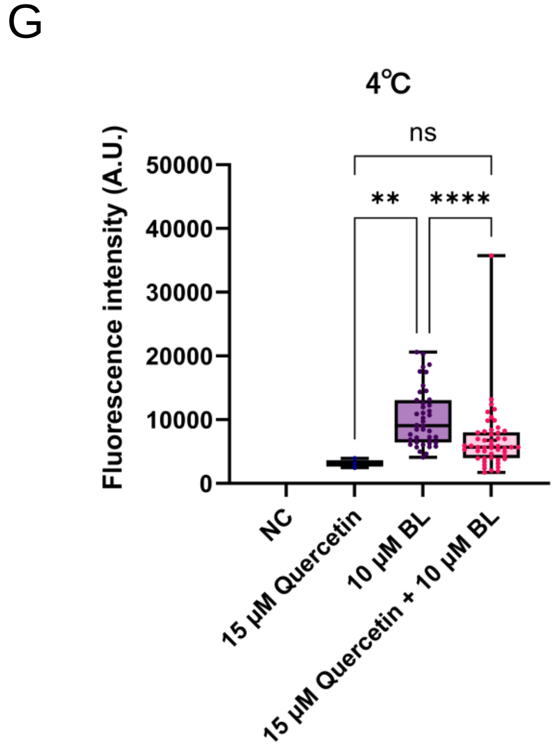


**Supplementary Fig. S3. Properties of pollen tubes incubated at very low temperature (4 ° C).**
A. Pollen tube germination rate at 4° C. Mean (SD), B. Pollen
B. tube length at 4° C. C– F. ROS staining of *in vitro* cultured pollen tubes without any compound (C), with 15 μM quercetin (D), 10 μM BL (E), and 15 μM quercetin and 10 μM BL (F), respectively. Pollen tubes were stained with CM-$H_2$DCFDA for 20 min. Scale bar, 50 μm. G. Quantification of ROS fluorescence intensity of pollen tube tip region (100 μm from the tip) at 4° C.

**Supplementary Data Set 1 (separate file).**

Normalized RNAseq cout data of wild type and *bri1-10* pistils, shown as reads per million mapped reads.

**Supplementary Data Set 2 (separate file).**

Downregulated genes in *bri1-10* pistils compared to those in wild type ((logFC < -1, FDR < 0.05, >1 rpm)
